# Potential role of Mercury pollutants in the success of Methicillin-Resistant *Staphylococcus aureus* USA300 in Latin America

**DOI:** 10.1101/2020.07.01.150961

**Authors:** C. A. Gustave, J.P. Rasigade, Patricia Martins-Simões, F. Couzon, Chloe Bourg, Anne Tristan, Frédéric Laurent, T. Wirth, F. Vandenesch

## Abstract

Community-acquired methicillin-resistant *Staphylococcus aureus* (CA-MRSA) lineage known as USA300-North American (NA) has become highly prevalent in North America whilst a USA300 variant known as USA300-LV, harboring a mercury resistance element (COMER), has become dominant in South America. We investigated whether mercury pollution, which is common in South America notably because of artisanal gold mining, may explain the local dominance pattern of USA300-LV. Density-based estimation of epidemic success in 250 genomes of the ST8 lineage revealed that the acquisition of COMER in USA300 progenitors increased success in South American countries but decreased success elsewhere. The fitness of USA300-LV was impaired *in vitro* compared with USA300-NA, but the addition of sub-inhibitory concentration of mercury provided a strong fitness advantage to USA300-LV and triggered an overexpression of major virulence factors. The success of USA300-LV in South America may result from low-level mercury exposure selecting resistant and virulent strains.

## Introduction

*Staphylococcus aureus* colonizes asymptomatically one third of human population and is also an important human pathogen responsible for a large diversity of both nosocomial and community-acquired infections. Until 1990s, methicillin-resistant *S. aureus* (MRSA) infections were essentially healthcare associated and were caused by a limited number of successful clones. Antibiotic-selective pressure present in health care setting was the prevailing hypothesis for the success of these clones in these environments whilst their impaired fitness explained their lack of epidemiological success in community setting (*1*). In the early 2000, infections caused by novel MRSA clones were reported in healthy individuals without known connections to healthcare institutions (*2, 3*). These community-acquired (CA)-MRSA emerged independently on three continents and comprised notably the sequence type 8 (ST8) SCC*mec*IVa (standing for staphylococcal chromosomal cassette encoding methicillin resistance gene of type IVa) pulsotype USA300 in the USA (abbreviated to “USA300” below), the ST80 SCC*mec*IV in Europe, North Africa and Middle-East (here referred to as “EU-ST80”), and the ST30 SCC*mec*IV in Oceania (*4*). Within a decade, the USA300 lineage become the most prevalent *S. aureus* lineage in the community in the US, and subsequently become also highly prevalent in healthcare setting (*5-8*). In the meantime, a variant of USA300 referred to as USA300 Latin variant (LV) became prevalent in South America, being the dominant CA-MRSA clone in Colombia, Ecuador and Venezuela, accounting for 10.6% of consecutive *S. aureus* isolates in hospital laboratories (*9-12*). The noticeable difference between USA300 strains from North America (USA300-NA) and USA300-LV are the presence in USA300-NA of a mobile element (ACME) encoding for factors promoting defenses against skin acidic pH and skin innate immune defenses (*13, 14*), and the presence in USA300-LV of a locus (COMER) encoding resistance to mercury (*11*). One of the human uses of mercury with the highest associated Hg emissions is its use for extracting gold in artisanal and small-scale gold mining (ASGM) (*15, 16*). Mercury emission from ASGM account for 37.1% of global anthropogenic mercury emissions to air in 2010, countries of Latin America being major contributors (e.g. Colombia 60.000 t/y, Bolivia 45.000 t/y, Peru 26.250 t/y, Brazil 22.500 t/y, Ecuador 17.500 t/y) whilst developed countries such as the USA had no such emission from ASGM (*15, 16*). Overall South America contributed to 12.5% (245 t/y) of global anthropogenic mercury emission whilst North America to solely 3.1% (60.7 t/y). The aim of this paper was to investigate whether mercury pollution may explain the success of USA 300-LV in South America, notably in the Andean countries where the high prevalence of USA300-LV (*9-12*) coincides with high environmental mercury pollution (*15*). Using the Timescaled Haplotypic Density (THD) method to examine patterns of epidemic success (*17*) in 250 genomes of the ST8 lineage, we confirmed that the acquisition of *mec*A, followed by COMER or ACME contributed to epidemic success of the two sub-lineages, and that COMER only increased success in South American countries. *In vitro* experiments in the presence of subinhibitory concentration of mercury revealed an overexpression of major virulence factors by USA300-LV and a fitness advantage of USA-LV over USA300-NA, thus providing a plausible model and selection drive explaining the success of USA300-LV in South America.

## Results

### The epidemic success of the ST8 lineage increases sequentially with acquisition of various mobile genetic elements

In a first attempt, we tried to confirm that the 137 strains (76 strains from the USA300-NA lineage and 61 strains from the USA300-LV lineage) correspond to measurably evolving populations. This was indeed the case with significant positive correlations **(Fig. S1 panel C)** between genetic distance and sampling time in both settings (USA300-NA and USA300-LV). The estimated short-term mutation rates corresponded to 4.64 x 10^−7^ (95% confidence interval of 3.04 x 10^−7^ to 6.28 x 10^−7^) substitutions per nucleotide site per year for the USA300-NA strains and to 6 x10^−7^ (95% confidence interval of 3.53 x 10^−7^ to 8.55 x 10^−7^) substitutions per nucleotide site per year for the USA300-LV strains. Those results are slightly dissimilar from previous reported rates based on all genomes SNPs (*18, 19*). However, this trend is not surprising because the present study focused on the sole housekeeping genes from the core-genomes, leading to a slower molecular clock rate. Maximum likelihood trees (GTR + gamma) were congruent with former tree topologies **(Fig. S2)** and show a fast diversification pattern.

The TMRCA of both lineages largely overlapped and date back to the early and mid 1980’s **(Fig. S1 panel A)**. This result is rather congruent with dating’s obtained by Planet and colleagues (*11*), as well as Straus and colleagues (*20*). We then generated a Bayesian skyline plot that estimated the pathogen’s demographic changes over time **(Fig. S1 panel B)**. Both, USA300-NA and USA-300LV displayed similar effective population size changes trough time with a first onset in the early 1990’s, a plateau at the beginning of the 21^st^ century, followed by a regular decline.

We compared the evolution of epidemic success in the two USA300 sublineages using time-scaled haplotypic density (THD) (*17*). The THD method uses pairwise genetic distances to estimate the epidemic success of each isolate in a collection, allowing comparisons of success based on various isolate characteristics. THD approaches have been successfully applied to investigate drivers of epidemic success in bacterial pathogens including *Staphylococci* (*17, 21-23*). Applied to 250 genomes of ST8 (combining those previously analyzed by (*20*) completed by 21 USA300-LV from our collection), the method revealed several important features: (i) the acquisition of the Panton-Valentine leucocidin gene by the ancestral ST8 clearly boosted the epidemic success of the lineage **(Fig. 1 panel A)**, (ii) the acquisition of *mec*A further increased the epidemic success, in accordance with the results obtained previously by coalescent-based demography approaches on this lineage **(Fig. 1 panel B)** (*24*), (iii) the acquisition of COMER or ACME further increased the epidemic success of the two sub-lineages, more strongly so for ACME than for COMER **(Fig. 1 panel C)**. Among USA300 isolates, defined here as having both the PVL and *mec*A determinants, the presence of COMER was associated with strongly decreased success indices in isolates collected outside of South America **(Fig. 1 panel D)**. We used linear regression to quantify to which extent the relationship of COMER with epidemic success depended on the geographic origin of USA300 isolates. Using the THD index as the response variable and COMER presence, isolation in a South American country and their interaction as predictors, both COMER and isolation in South America were independently associated with a decreased success (coefficients -0.52 (95% CI, -0.75 to -0.28) and -0.54 (−0.85 to -0.23), respectively), however success strongly increased when the COMER-positive isolate was isolated in South America (interaction term coefficient, +0.69 (0.29 to 1.08)).

**Figure 1.**
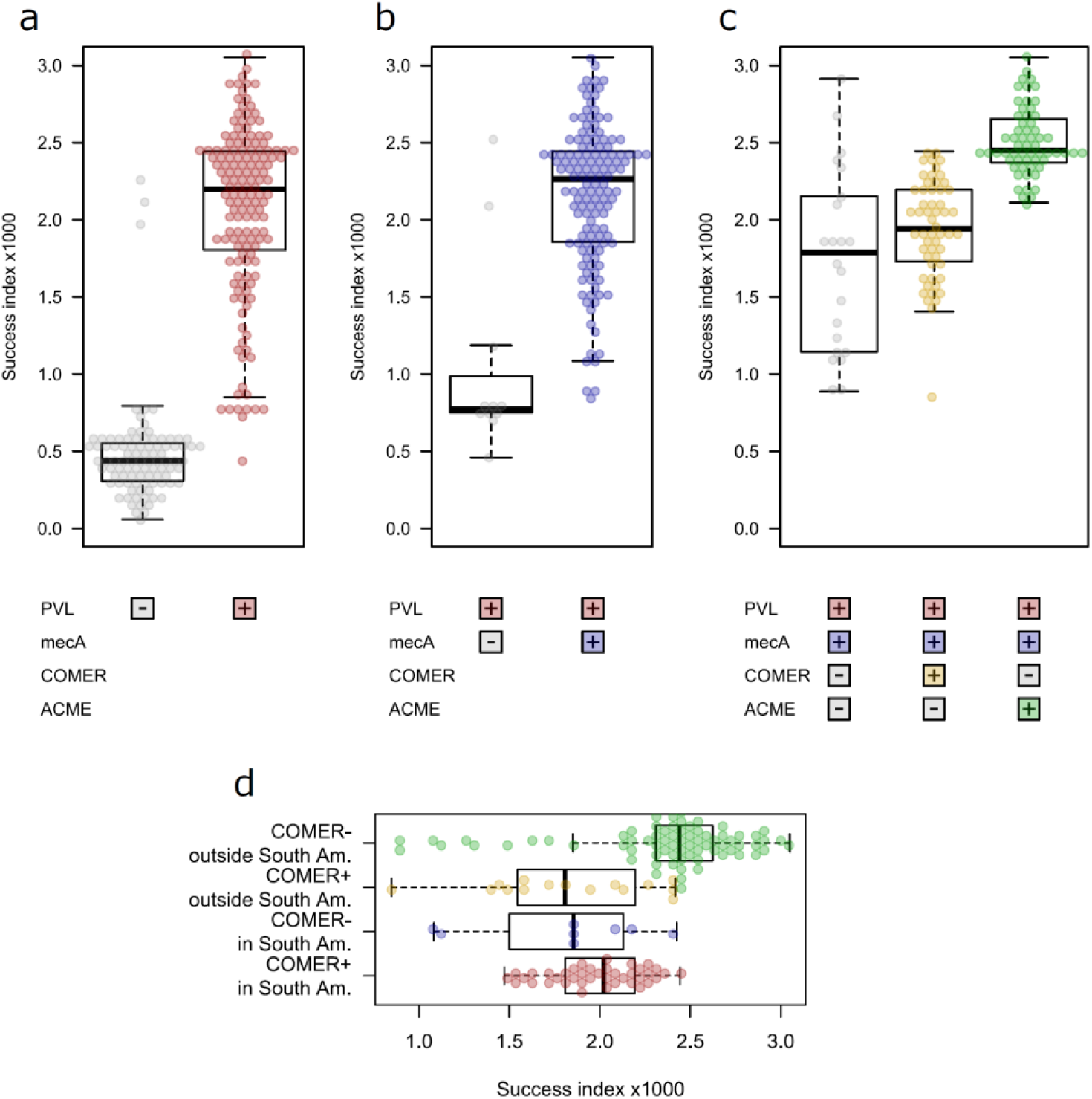
Density-based estimates of epidemic success in 250 *S. aureus* ST8 genomes vary with the presence of mobile genetic elements. **(A-C).** Evolution of timescaled haplotypic density (THD) indices with the sequential acquisition of the phage encoding the Panton-Valentine leukocidin (PVL) **(A)**; of the methicillin-resistance determinant *mec*A in PVL-positive isolates, leading to the emergence of the USA300 lineages **(B)**; and of either ACME or COMER **(C)**, leading to the emergence of the USA300-NA and USA300-LV sublineages, respectively. (**D)**, COMER acquisition in USA300 (PVL-positive and *mec*A-positive ST8 *S. aureus*) is associated with lower success in non-South American countries but with higher success in South American countries.

### The USA300-LV display an overall fitness impairment in comparison to USA300-NA

We conducted *in vitro* experiments to test whether the epidemiological link between mercury resistance in USA300-LV and mercury pollution in South America was associated with a measurable phenotype. We first compared the fitness of USA300 strains representatives of the different steps of evolution of this lineage, i.e. before and after the acquisition of methicillin resistance (ancestral ST8 MSSA and basal USA300 MRSA respectively), followed by acquisition of ACME for the NA variant (derived USA300-NA MRSA, including early-derived non-multi-resistant strains and late-derived multi-resistant strains), or the acquisition of COMER for the LV variant (derived USA300-LV MRSA) **(Table 1) (Fig. 2)**. As shown previously (*25*), doubling time measurement revealed that the acquisition of methicillin resistance (SCC*mec*) was associated with impaired fitness that was totally compensated by the acquisition of ACME in early-derived USA300 NA **(Fig. 3)**; however, the fitness cost of the acquisition of other resistances (such as FQ resistance) in late-derived USA300 impaired the fitness cost to a higher level **(Fig. 3)**. In parallel, the acquisition of COMER in USA300-LV was associated with increased doubling time at an intermediate level between those of early and late-derived USA300-NA strain **(Fig. 3)**.

**Table 1.** Bacterial strains with HgCl_2_ MICs. HgCl_2_ minimum inhibitory concentrations (MIC) determined by microdilution method for each strain identified in column 1 and including major discrimination determinants for each sublineages in column 2.

| Strain designation | Relevant characteristics | HgCl<sub>2</sub> MIC (mg/L) |
| --- | --- | --- |
| Ancestral ST8 ST20172183 | *mecA*(-) ACME(-) COMER (-) | 0.57 |
| Basal USA300 ST20172178 | *mecA*+ ACME (-) COMER (-) | 0.57 |
| USA300-LV 1 ST20172176 | *mecA*+ ACME(-) COMER+ | **6.8** |
| USA300-LV 2 ST20172184 | *mecA*+ ACME(-) COMER+ | **6.8** |
| USA300-LV 3 HT20030343 | *mecA*+ ACME(-) COMER+ | **6.8** |
| USA300-NA 1 ST20120558 | *mecA*+ ACME+ COMER (-) | 0.57 |
| USA300-NA 2 ST20111414 | *mecA*+ ACME+ COMER (-) | 0.57 |

**Figure 2.**
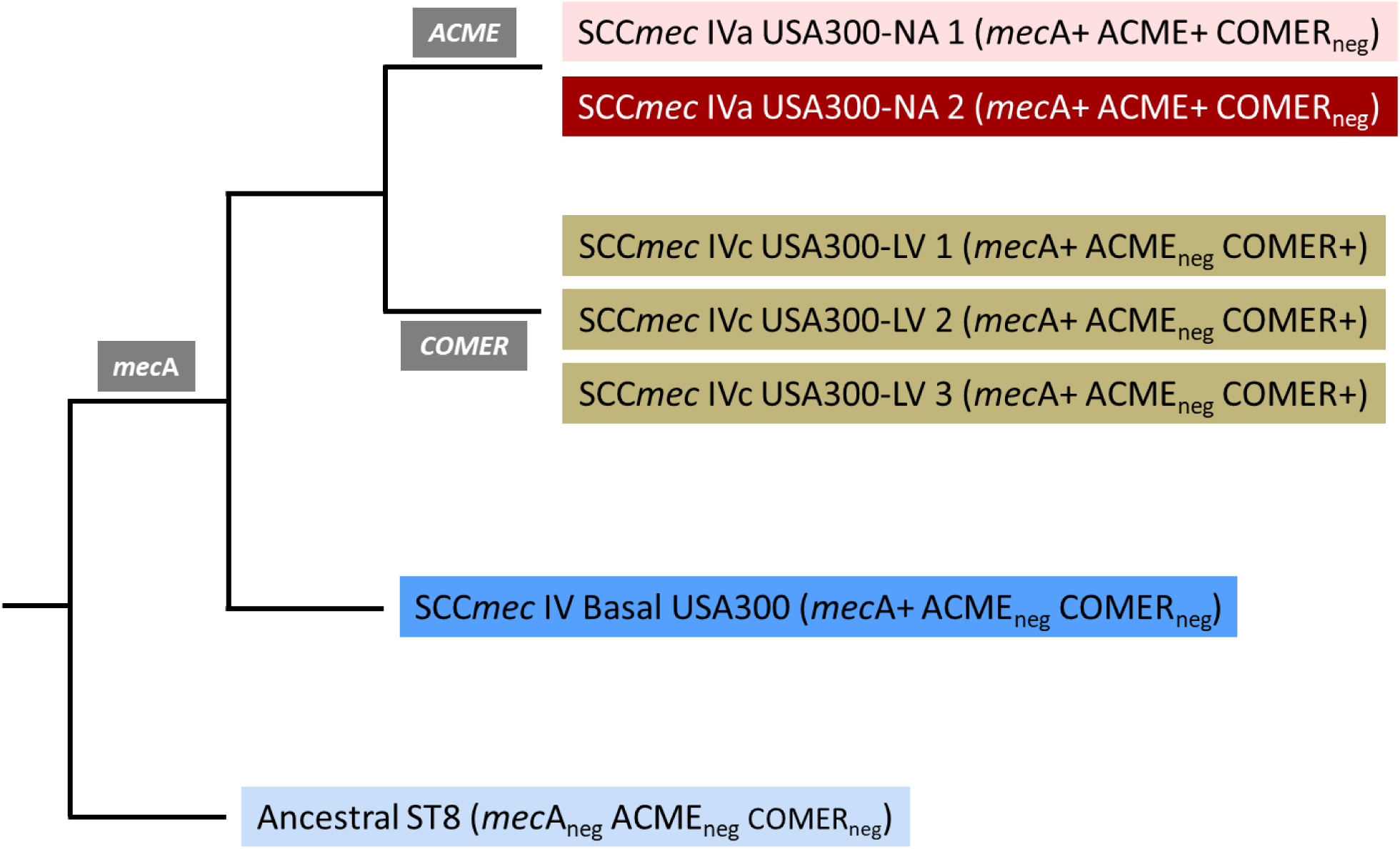
Schematic representation of the strains relative positions to evolutionary key-points of USA300 NA and LV lineages. Denomination of each strain includes major discrimination parameters for each sublineages indicated by the color code used in subsequent figures.

**Figure 3.**
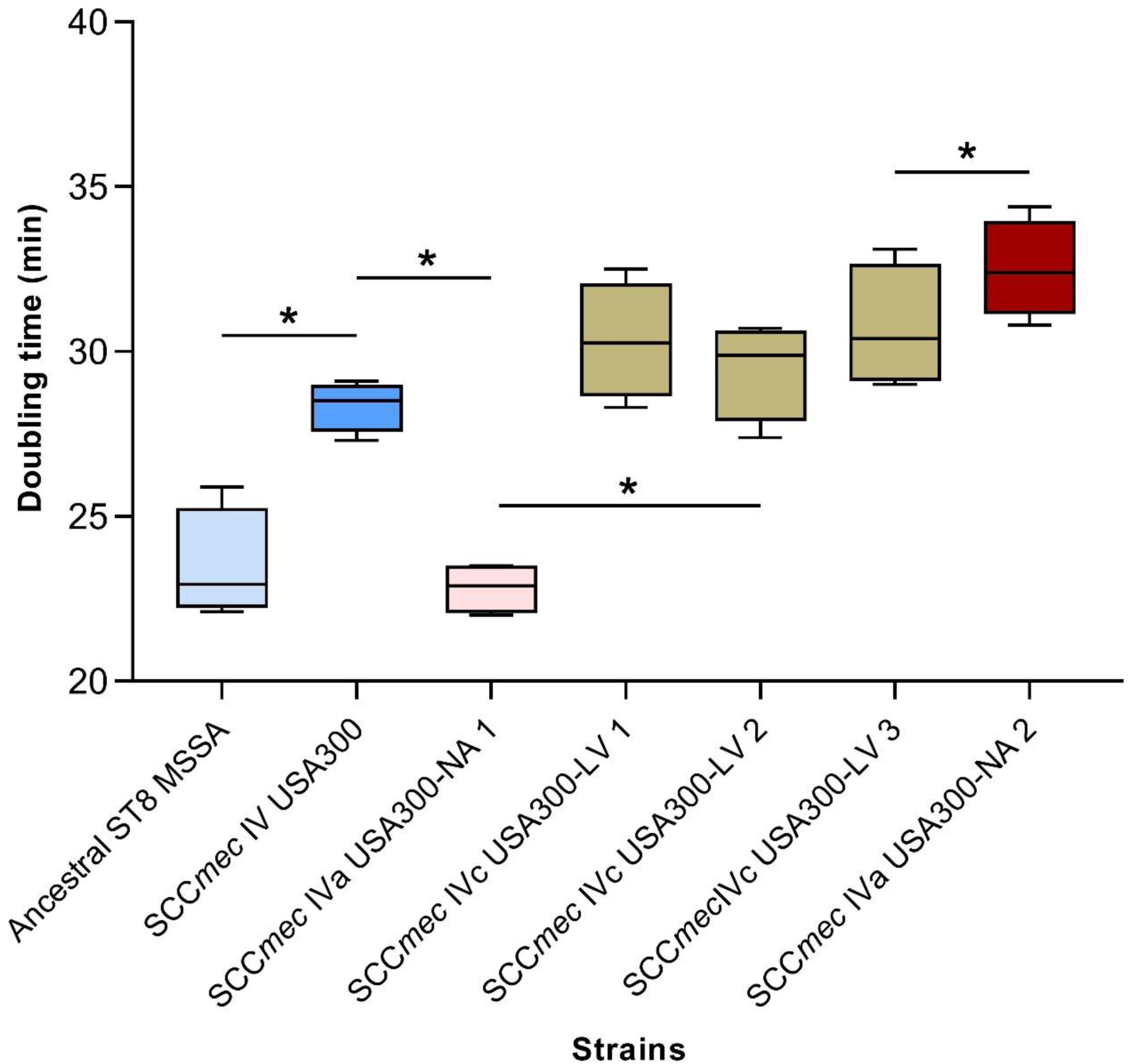
Doubling times of ST8 lineage strains. ST8 isolates were cultured in BHI incubated on 96-wells plates for 24 hours at 37°C with continuous optical density monitoring at 600nm (Tecan Infinite® 200 PRO). Doubling times were calculated by graphical method after Log transformation of data from the exponential growth phase. The color codes for each strain correspond to those in figure 2 (*: *P* = 0.029). Experiments were performed on three independent series (biological replicates), and optical densities were measured on three wells for each strain (technical replicates).

Competitive fitness between the different strains showed that in all cases, the strains with shorter doubling time outcompeted those with longer doubling time, and thus the LV was totally outcompeted by the ancestral ST8 **(Fig. 4 and Fig. S3)**.

**Figure 4.**
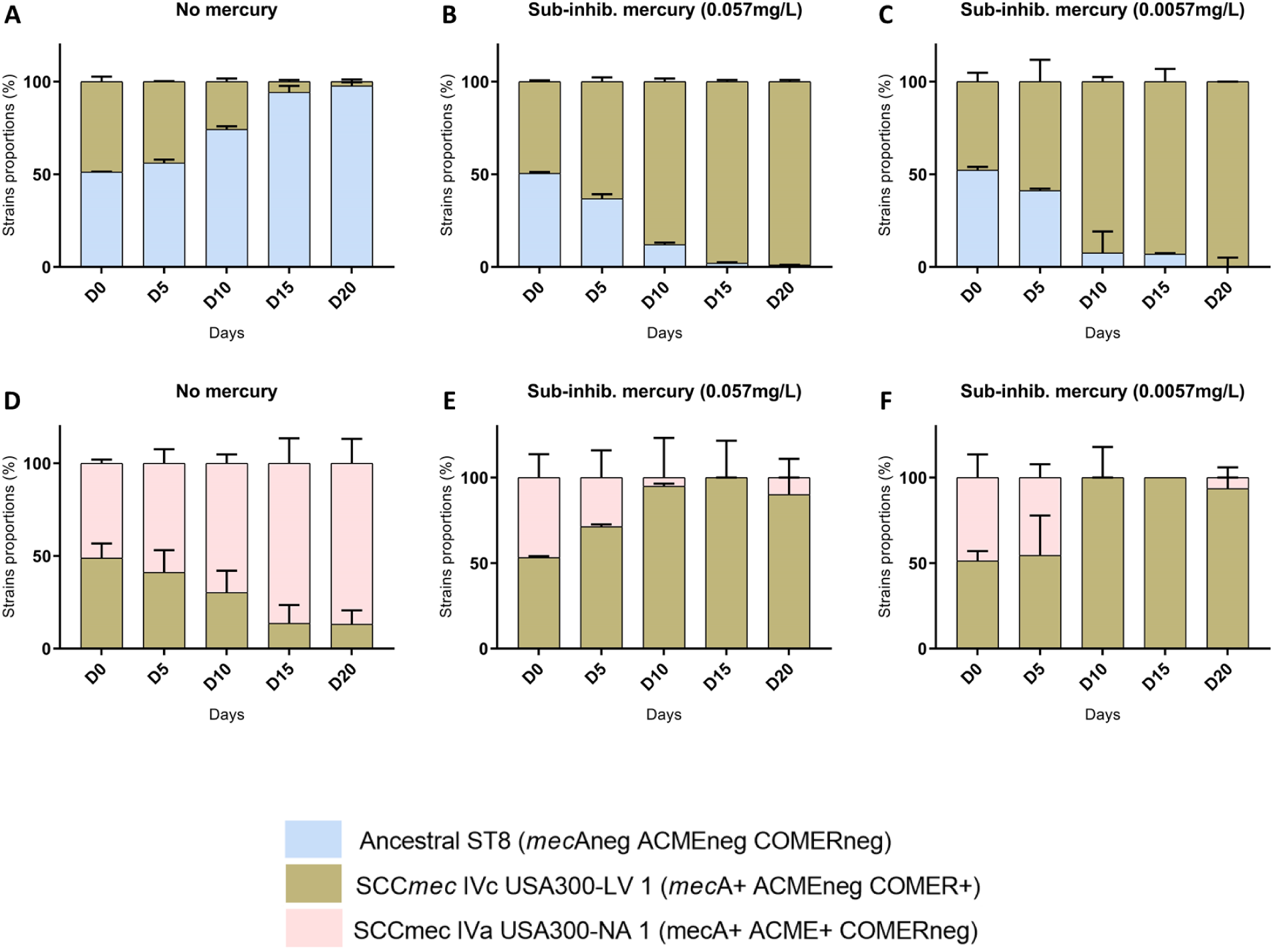
Impact of ACME, COMER and mercury resistance on competitive fitness of USA300 NA and LV variants. **(A)** USA300-LV (COMER-positive) strains and ancestral ST8 MSSA were co-cultivated for 20 days in BHI without antibiotics or **(B)** containing HgCl_2_ at ^1^/_10_ of HgCl_2_ MIC of the susceptible strain (0.057 mg/L), or **(C)** ^1^*/*_100_ MIC (0.0057 mg/L) with daily subculture in fresh medium. **(D)** USA300-NA (ACME-positive) and USA300-LV (COMER-positive) strains, both MRSA were co-cultivated for 20 days in BHI without antibiotics or **(E)** containing HgCl_2_ at ^1^/_10_ of HgCl_2_ MIC of the susceptible strain (0.057 mg/L), or **(F)** ^1^*/*_100_ MIC (0.0057 mg/L) with daily subculture in fresh medium. The proportion of each strain was monitored at day 0, 6, 10, 15 and 20 with differential colony count based on selective agar inoculated with a calibrated amount of competitive mix. Competitive cultures were performed on three independent series (biological replicates), and each colony count was repeated three times (technical replicates).

These results, notably the strong impairment of USA300-LV toward early USA300-NA cannot recapitulate the observed field epidemiology where USA300-LV was shown to be the main CA-MRSA in the Andean countries at the beginning of the 21^st^ century when the possible USA300-NA competitor was still the early-derived USA300-NA (FQ susceptible) (*11, 25*). In addition, these results are counterintuitive to the above results of THD showing increased success of both COMER and ACME sub-lineages. We thus questioned the role of mercury-resistance, encoded in the COMER element, in the success of the USA300-LV in the Andean countries, taking into account the fact that mercury pollution from mining and ore processing is particularly prevalent in these countries (*15*).

### Mercury resistance confers a fitness advantage to USA300LV in the presence of Mercury

HgCl_2_ MICs were determined in COMER-positive and-negative strains, revealing that the level of mercury resistance was ten-fold higher in COMER-positive strains as compared to the COMER-negative ancestral ST8, basal USA300 or USA300-NA **(Table 1)**. We then performed competitive fitness experiment in the presence of HgCl_2_ at various concentrations ranging from 1:10 to 1:1000 of susceptible strain MIC. After 20 days of coculture, the COMER-positive strains outcompeted all COMER-negative strains, notably the USA300-NA early- and late-derived (**Fig. 4 panel E & F**, and **Fig. S4**). This effect was observed at 1:10 and 1:100 of susceptible strain MIC (0.057 mg/L and 0.0057 mg/L, respectively). These results reveal that low-level exposure to HgCl_2_ – a condition that could typically occur on the skin of workers in artisanal gold mining (*15*) provide a selective advantage to the USA300-LV over the USA300-NA whether early or late-derived.

### Mercury exposure induces the expression of virulence factors

Overexpression of core-genome encoded virulence factors has been proposed to contribute to the expansion of CA-MRSA (*26*). Given the known global transcriptomic and proteomic impact of mercury on bacteria (*27, 28*) and the mercury resistance of USA300-LV, we wondered whether mercury could be a factor of increased expression of virulence factors in this lineage. To this end, RT-PCRs targeting virulence factors of the core- (α-toxin, PSMα, γ-toxin) and accessory-genome (LukS-PV), as well as the major regulator of *S. aureus* virulence (agr-RNAIII), were performed after *in vitro* post-exponential growth as previously described (*26*). These experiments performed without HgCl_2_ and upon exposure to sub-inhibitory concentration of HgCl_2_ (^1^/_10_ of HgCl_2_ MIC) revealed a constant increase of virulence factors expression (7-11 fold for *hla*, 4-38 fold for *hlg*C, 6-12 fold for *luk*S-PV, 6-7 fold for *psm*α) and of the regulatory *agr*-RNAIII (3.5-13 fold) upon exposure of all three USA300-LV strains to HgCl_2_. In contrast the ancestral ST8 MSSA, the basal USA300 MRSA as well as the two derived USA300 NA showed either a down-regulation or no variation of virulence factor expression in the presence of mercury **(Fig. 5)**.

**Figure 5.**
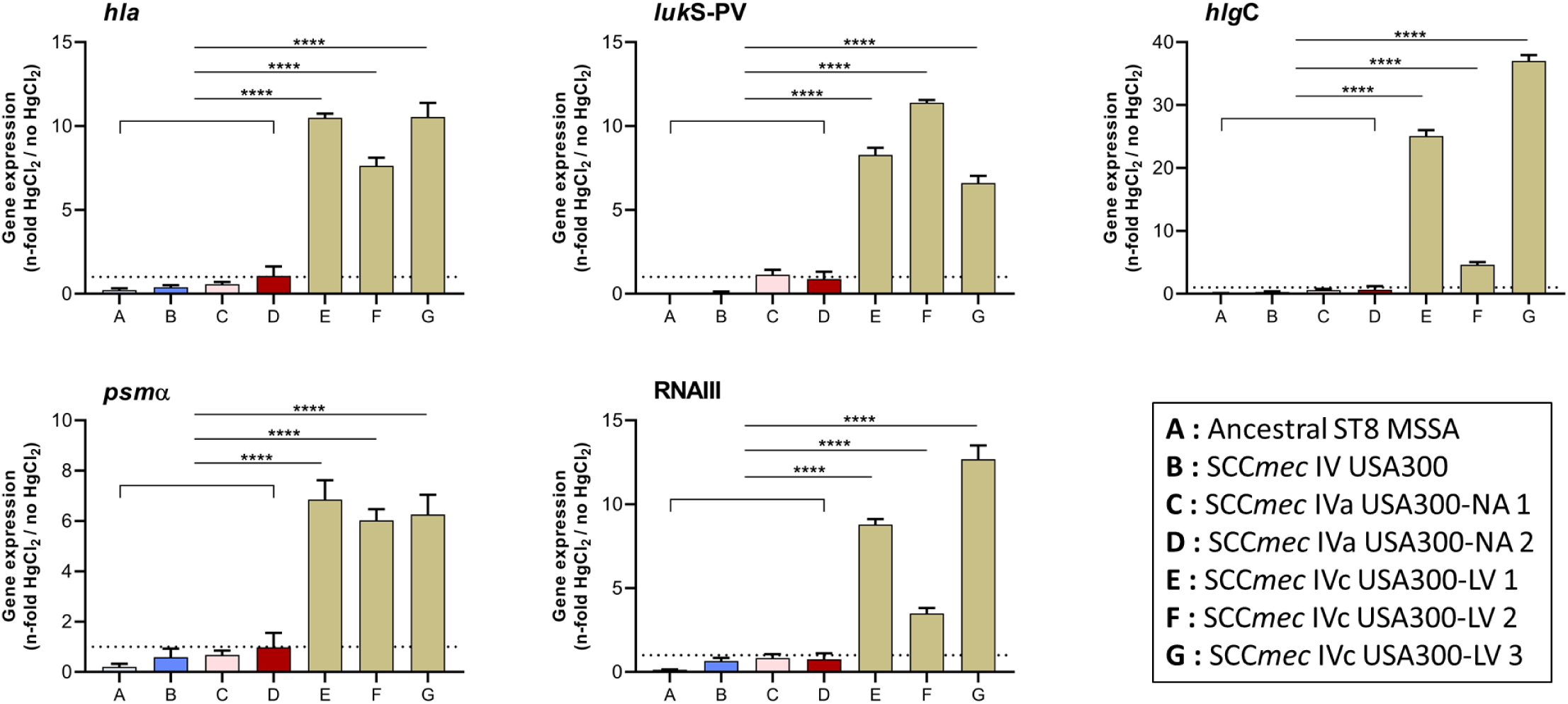
Mercury enhance expression of virulence related genes in USA300LV strains ST8 strains. Expression of virulence factor- and regulatory-genes were assessed by qRT-PCR among ST isolates exposed to ^1^/_10_ of HgCl_2_ MIC (table 1). Results are expressed as fold change in comparison to the same strain without HgCl_2_ exposure. Experiments were performed on three independent series (biological replicates), and three RNA quantifications were done for each RNA sample (technical replicates).

## Discussion

Both THD and coalescent-based demography revealed that the two USA300 sublineages increased their epidemic success and their effective population size upon acquisition of *mec*A. These results could sound counterintuitive with respect to the increased biological cost associated with the acquisition of the antibiotic resistance locus but are in accordance with previous results showing that this biological cost is totally reversed in the presence of trace amount of antibiotics that can be present in the environment (*25*). Similarly, whilst ACME was associated with a gain in fitness, COMER was associated with an increased doubling time and a decreased competitive fitness when challenged by ancestral ST8 MSSA or USA300-NA strains **(Fig. 3 and 4 panel A)**; however, sub-inhibitory concentrations of HgCl_2_ totally reversed the fitness cost of COMER, allowing USA300-LV to outcompete all other sub-lineages **(Fig. 4, Fig. S3 and Fig. S4)**. Noticeably THD revealed that the acquisition of PVL was associated with an increased epidemic success of the ST8 lineage. The role of PVL as a factor contributing to the success of the historical major CA-MRSA lineages (USA300, European ST80, ST30) (*3*) has been largely debated (*3, 29, 30*). As a factor associated with skin and soft tissues infections (*31*), and particularly primary skin infections such as folliculitis and furuncles (*32-34*), PVL could indirectly promote dissemination between humans by increasing *S. aureus* bacterial load associated with cutaneous infections and thus enhances the epidemic success of strains harbouring PVL. This suggest that acquisition of the PVL which is carried by a bacteriophage, should enhance the epidemic success on any lineages whatever MSSA or MRSA. Unfortunately, this is unlikely to be provable as there is no comparable situation to the ST8 lineage in which isolates before and after the acquisition of PVL can be studied at the genomic level using coalescent-based demography and THD approaches. A speculative example is the case of the methicillin-sensitive phage type 80/81 clone of the CC30 lineage, in the 1950s and 1960s, which spread from hospitals causing a significant disease burden in the community and was characterized by resistance to penicillin and production of PVL (*35-38*). According to McAdam *et al*. (*39*) PVL have been maintained in CC30 clades since a likely acquisition event that occurred about 137 years ago, these data being consistent with a central role for PVL in the success of some community-associated *S. aureus* clones (*29, 39*). In a similar reasoning, linking cutaneous infections to clonal expansion in the community, the success of CA-MRSA has been partly attributed to increased expression of core-genome encoded virulence factors (*26*), enhancing cutaneous infection rate and thus human-to-human transmission by skin contact as observed in prisons, sport team or men-having-sex-with-men (*11*). Here we showed that expression of several major virulence determinant (including toxin and regulatory genes) is specifically up-regulated in USA300-LV in presence of sub-inhibitory concentrations of mercury. While the mechanism is still not known, such concentrations of mercury can be present on the skin of workers employed in artisanal gold mining which is well developed in Andean countries and where HgCl_2_ is used in the final amalgamation steps of ore processing. As noticed by the Pure Earth Blacksmith Institute, due to a lack of awareness, as well as lack of environmental, health, and safety regulations in these small mining industries, miners are often exposed to dangerous levels of toxic materials by inhalation and ingestion (*40*). These conditions provide a selective advantage of resistant strains (USA300-LV) over susceptible one (USA300-NA). The *mer* operon enables wide-spectrum resistance to mercury through a binary system composed of i) *mer*A (mercuric reductase needed for reduction-based mercury detoxification), ii) *mer*B (organomercurial lyase that cleaves the carbon-mercury bond of organomercurials for subsequent detoxification by the mercuric reductase) (*41*). This system is under *mer*R regulation. This *mer* operon also includes a sequence coding for a putative cytochrome C biosynthesis protein. Its position within the *mer* operon might suggest a coregulation of the *mer*AB system and respiratory adaptation in mercury enriched environment. Altogether this mercury resistance confers a selective advantage of USA300-LV strains, not only because of direct resistance capacity toward the toxic effect of HgCl_2_, but also because at such tolerable concentrations of HgCl_2_, the strains are likely more virulent and thus more prone to produce skin infections which favor human-to-human transmission. Overall anthropogenic activities leading to environmental pollution may have driven the expansion of USA300-LV in South America and noticeably in the Andean countries. Worth mentioning, the TMRCA of the USA300-LV clade is in frame with an historical peak in gold price *(42)*, an external factor that might boost the motivation for excavating activities. In contrast, in countries where exposure to mercury is less common, such as North America (*16*), the biological cost of the COMER encoding mercury resistance is counterselective, allowing the USA300-NA variant to outcompete the USA300-LV. USA300-LV is not the only MRSA lineage present in South America; of note some isolates belonging to ST72, ST88, and ST97 harbor some fragments of the COMER island (notably *mer*R, *mer*A and *mer*B) (*12*), supporting the hypothesis that mercury resistance provide a selective advantage in countries where environmental exposure to mercury is discernible (*15*).

## Materials and Methods

### Bacterial strains and genomes

Genomic data from 250 strains belonging to ST8, CA-MRSA USA300-NA and USA300-LV (also denominated SAE for South American Epidemic) were combined **(Table S1)**. This included genomic data of 223 isolates from the study by (*20*), 19 isolates from the study by (*12*) and 8 USA300-LV isolates from the present study. These 8 strains were from 8 sporadic cases (infection or carriage) reported to the French National Reference Center for Staphylococci (NRC Staphylococci) during the past 15 years. Sequencing of those 8 isolates was performed using Illumina MiSeq (2×300 bp) or NextSeq (2×150 bp) sequencers, with a coverage of >30×. Libraries were constructed with the Illumina Nextera XT kit, as recommended by the manufacturer. Sequence reads from previously described USA300 strains were downloaded from the European Nucleotide Archive website (http://www.ebi.ac.uk/ena) or the NCBI assembly (https://www.ncbi.nlm.nih.gov/assembly) website. SNP prediction was performed with Snippy (https://github.com/tseemann/snippy) and indications of putative horizontal gene transfer events were defined as a unique SNPs cluster (≥3) and purged from the alignment. For functional studies (MICs, fitness, virulence), isolates were selected at various temporal steps of their inferred population dynamics. For the CA-MRSA USA300 lineages ancestors, 2 clinical isolates were included **(Table 1 & Fig. 2)**: (i) one ancestral strain of the ST8 lineage, susceptible to methicillin and lacking ACME or COMER (Ancestral ST8 MSSA), (ii) one strain corresponding to the most recent common ancestor of the USA300-NA and LV clones, being resistant to methicillin but lacking the ACME or COMER sequence (Basal USA300 MRSA). For the CA-MRSA USA300-NA lineage, 2 clinical isolates were included **(Table 1 & Fig. 2)**: (i) 1 strain from the early expansion phase characterized by ACME and SCC*mec* acquisition (USA300-NA 1 MRSA), (ii) 1 strain from the most recent evolutionary phase subsequent to fluoroquinolones resistance acquisition (USA300-NA 2 MRSA). For the CA-MRSA USA300-LV lineage, 3 clinical isolates were included **(Table 1 & Fig. 2)**, all lacking ACME but carrying COMER sequence (USA300-LV 1, 2 & 3).

### Phylogenetic and Coalescent based analysis

For the phylogenetic reconstruction a subset of 137 samples (76 strains from the USA300-NA lineage and 61 strains from the USA300-LV lineage) were selected according to their phylogenetic position in the USA300-NA clade or LV clade as previously published (*20*) SNP prediction was performed for each set: NA and LV using as reference a ST8 MSSA PVL+ (ancestral to both USA300 SAE and NAE clades, strain ID 800903 in (*20*). Since no traces of recombination were detected (*24*), the phylogenetic relationships were directly reconstructed by the ML approach implemented in PhyML 3.0 (*43*). The robustness of the ML tree topology was assessed with bootstrapping analyses of 1,000 pseudoreplicated data sets. Phylogenies were rooted with strain Germany_2_2008.

Evolutionary rates and tree topologies were analyzed with the generalized time reversible (GTR) (*44*) and Hasegawa-Kishino-Yano (HKY) (*45*) substitution models with gamma distributed among-site rate variation with four rate categories (Γ4). We tested both a strict molecular clock (which assumes the same evolutionary rate for all of the branches of the tree) and a relaxed clock that allows different rates among the branches. Constant-size, logistic, exponentially growing coalescent models were used. We also considered the Bayesian skyline plot model (*46*), based on a general, nonparametric prior that enforces no particular demographic history. We used a piecewise linear skyline model with 10 groups and then compared the marginal likelihood of each model with Bayes factors estimated in Tracer 1.7.1. Bayes factors represent the ratio of the marginal likelihood of the models being compared. Approximate marginal likelihoods for each coalescent model were calculated via importance sampling (1,000 bootstrap replications) with the harmonic mean of the sampled likelihoods. A ratio between 3 and 10 indicates moderate support of the idea that one model fits the data better than another, whereas values of >10 indicate strong support. For each analysis, two independent runs of 100 million steps were performed and the chain was sampled every 10,000th generation. Examination of the Markov chain Monte Carlo (MCMC) samples with Tracer 1.7.1 indicated convergence and adequate mixing of the Markov chains, with effective sample sizes for each parameter in the hundreds or thousands. The first 10% of each chain was discarded as burn-in. We found the maximum clade credibility topology with TreeAnnotator 1.10.4, and we reconstructed the Bayesian skyline plot with Tracer 1.7.1. The relaxed clock models provided a better fit to the data (Bayes factor, > 11) and under the different models tested, the Bayesian skyline model provided a (marginally) better fit, overall.

### Timescaled Haplotypic Density estimation

For the SNP distance matrix used in the THD model, the full set of 250 genomes was used. The reference strain used for the reads alignment was strain TCH1516 (NC_010079.1), as used previously by Strauß *et al.* (*20*). Briefly, the THD approach uses kernel density estimation in the space of genetic distances to assign a density estimate to each haplotype (isolate) in a collection. Such density estimates where shown both empirically and in simulations to correlate with the epidemic success in the ancestry of the isolate (*17, 23*). THD computations were conducted using R 3.6.1 with the *thd* package, available at github.com/rasigadelab/thd. The matrix of pairwise SNP distances was used as input; the THD parameters were an effective genome size (number of DNA base pairs effectively used in distance computations) of 3×10^6^ bp; a per-nucleotide mutation rate of 1×10^−6^ per year; and a timescale of 20 years, consistent with the recent expansion of the USA300 lineage.

### HgCl_2_ MIC determination

minimal inhibitory concentration (MIC) of HgCl_2_ were assessed by broth dilution. Strains were first grown on Columbia with sheep-blood agar (bioMérieux, France) overnight (ambient air, 35°C). For each strain, 3 to 5 colonies were transferred in Mueller-Hinton broth (Thermo Scientific Oxoid) and adjusted to an optical density at 600nm (OD) of 0.5. Suspension were then diluted to 1/100^th^. 100 µL of these suspensions were added to either 100µL of Mueller-Hinton broth, or 100 µL of HgCl_2_ Mueller-Hinton solutions (Fisher Chemical) to final HgCl_2_ concentrations of 6.8 – 1.7 – 0.56 – 0.17 – 0.057 – 0.017 – 0.006 mg/L in a final volume of 200µL. Suspensions were distributed in triplicate in 96-well round bottom plates (Greiner). Experiment was repeated on three different days (biological replicates).

### Crude doubling time

Isolates growth curves were determined from Brain Heart Infusion broth (BHI) cultures incubated in 96-well flat bottom plates (Greiner) for 24 h at 37 °C with continuous optical density monitoring at 600 nm (Tecan Infinite® 200 PRO). Each strain was inoculated in three independent wells (technical replicate), and the experiment was repeated on three different days (biological replicate). Doubling times were calculated by graphical method with the log-transformed optical density data of the exponential growth phase.

### Competitive fitness

Each strain to be tested in a competitive pair was adjusted to an OD600 nm of 1, then three mL of a 1/100^th^ dilution in BHI of each strain was mixed in a glass tube. For experiments involving HgCl_2,_ the later was added at final concentrations corresponding to 1/10^th^ to 1/100^th^ of the susceptible strain’s MIC. Tubes were incubated at 35 °C in aerobic atmosphere under agitation (200 r.p.m.) for 22±2 h, and 50μL were transferred daily for 21 days to a fresh tube containing three mL of BHI. The proportion of each strain in the competitive mix was monitored at day 0, 6, 10, 15 and 20 with differential colony count based on selective agar inoculated with a calibrated amount of competitive mix (Spiral System® - Interscience) followed by aerobic incubation for 24 h at 37 °C. For MSSA vs MRSA pairs, we used the ChromAgar® medium (i2A, France) allowing for growth of both strains (total count) and the ChromID-MRSA® medium (bioMérieux) for MRSA colony count. For MRSA strains couples (USA300-LV/USA300-NA, or USA300-LV/Basal USA300), differential colony count based on BHI agar without and with 1.14 mg/L HgCl2 was used. Continuous competitive cultures were performed on three independent series (biological triplicates); each colony count was performed on three technical triplicates. For all strain pairs tested, one of the strains was eventually reduced to a trace level, so no statistical test was required for strain proportion comparisons.

### RNA extraction from *S. aureus*

Brain Heart Infusion (BHI) broth was inoculated with an overnight culture to an initial OD600 nm of 0.05 and grown up on aerated Erlenmeyer flask to the end of exponential phase (6 h) at 37 °C under agitation (200 r.p.m.). One millilitre of bacterial suspension was harvested, and concentration adjusted to an OD600 nm of 1.5. Bacteria were washed in 10mM Tris-buffer and treated with lysostaphin and β-mercaptoethanol. RNAs were extracted with the RNeasy Plus Mini Kit® (Qiagen), quantified by spectrophotometry and stored at −80 °C. This process was repeated on three different days for biological replicates.

### RNA quantification by real-time PCR

A random-primers based reverse transcription of 1 μg of RNA was performed with the A3500 Reverse Transcription System Kit (Promega), followed by quantitative real-time PCR on cDNA using the FastStart Essential DNA Green Master kit (Roche) and the LightCycler® Nano (Roche). As previously described (*26*), we targeted five virulence genes (RNAIII, *luk*S-PV, *hla, hlg*C and *psm*α) and the house-keeping gene *gyr*B for normalization. Gene expression levels were compared between our clinical isolates; levels were expressed as n-fold differences relative to the most basal isolate of the strain set. These qRT-PCR’s were performed as technical triplicates (three RNA quantification per RNA sample), on RNA obtained from three biological replicates (three independent cultures and extractions per strain).

### Statistical analysis

Results are mostly presented as boxplots in figures and expressed as medians with 95% confidence interval (CI). Results from figure 3 are presented in boxplots and expressed as medians with min/max range obtained from three technical replicates and three independent biological replicates. In figures 4, S3 and S4, strains proportions in competitive pairs are presented as stacked histograms. All statistical comparisons were performed through non-parametric Mann–Whitney test (P-value <0.05). For competition experiments in which one strain was eventually reduced at undetectable level, no statistical test was required for proportion comparisons. All data were processed with Graphpad® PRISM 7.

### Data deposition

The data reported in this paper have been deposited in the European Nucleotide Archive (accession no. PRJEB38071).

## Supplementary Materials

Embedded at the end of this file:

**Figure S1. USA300 measurably evolving population dimension and demogenetics.**

**Figure S2. Evolutionary relationships of the selected strains belonging to the two USA300 sublineages.**

**Figure S3. Impact of COMER and mercury resistance on competitive fitness of USA300 ancestral strains and LV variants.**

**Figure S4. Impact of ACME, COMER and mercury resistance on competitive fitness of USA300 NA strains and LV variants.**

**Table S1. Combined genomic data from 250 strains belonging to ST8, CA-MRSA USA300-NA and USA300-LV.**

## General

We thank Sylvère Bastien, Michèle Bès, Laure Cazimajou, Typhanie Le Hir, and Karen Moreau for valuable technical and/or intelectual contribution.

## Author contributions

CAG designed the study, performed the experiments, interpreted the data, revised the manuscript. JPR and TW performed the experiments, interpreted the data, wrote the manuscript. PMS performed the experiments, interpreted the data, revised the manuscript. CB and FC performed the experiments and revised the manuscript. AT and FL contributed to the design of the study and revised the manuscript. FV designed the study, interpreted the data, wrote the manuscript.

## Supplementary Materials

**Figure S1.**
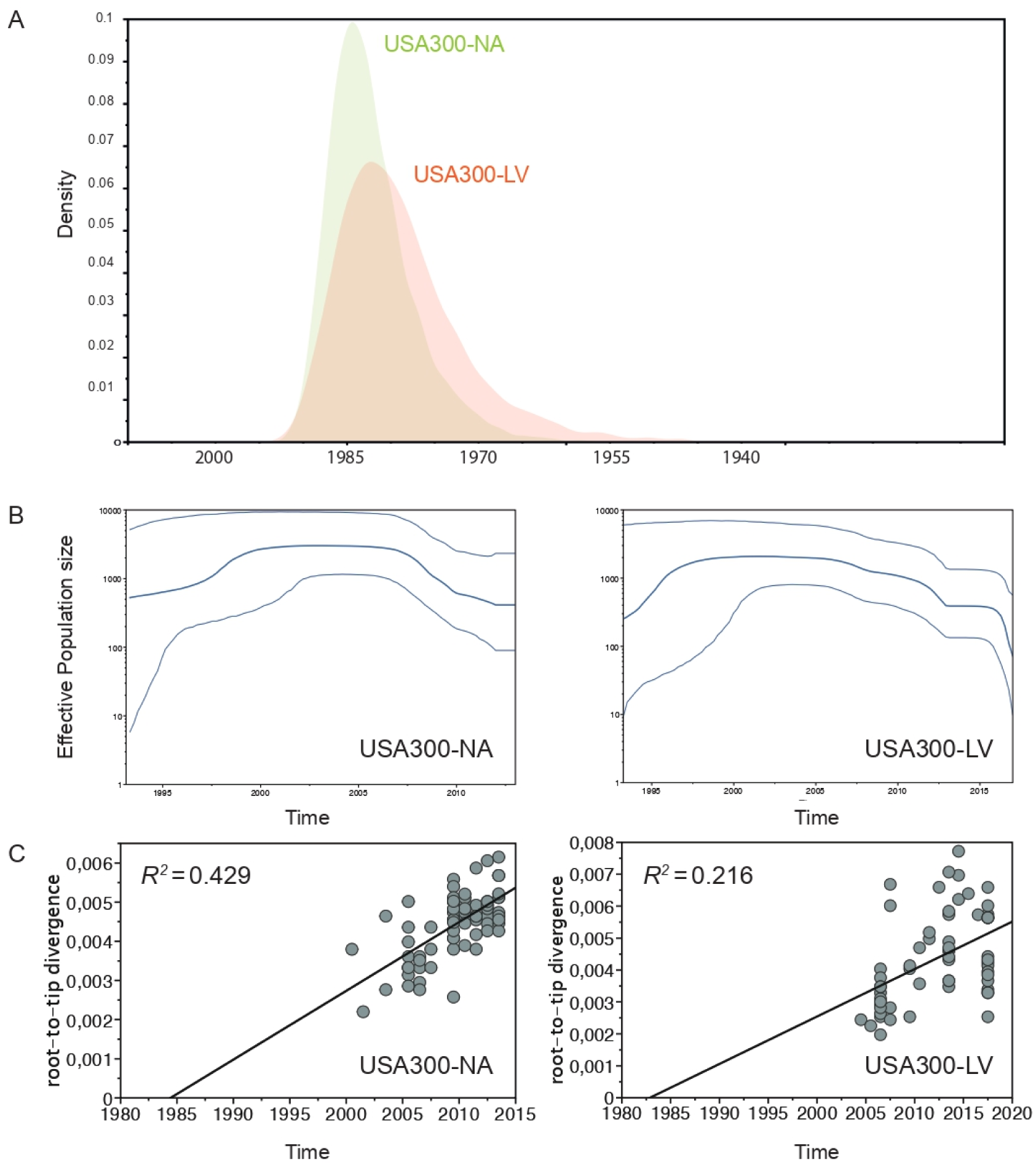
USA300 measurably evolving population dimension and demogenetics. **(A)**, Time to the most recent common ancestor (TMRCA) of the USA300-NA and USA-300-LV lineages. **(B)**, Bayesian skyline plots indicating population size changes in the two sublineages with a relaxed molecular clock. The fin lines represent the 95% confidence interval. **(C)**, Root-to-tip regression analyses. Plots of the root-to-tip genetic distance against sampling time are shown for the two sublineages phylogenies. The generated trees correspond to maximum-likelihood trees (GTR + gamma). Sample dates are given as years before the present.

**Figure S2.**
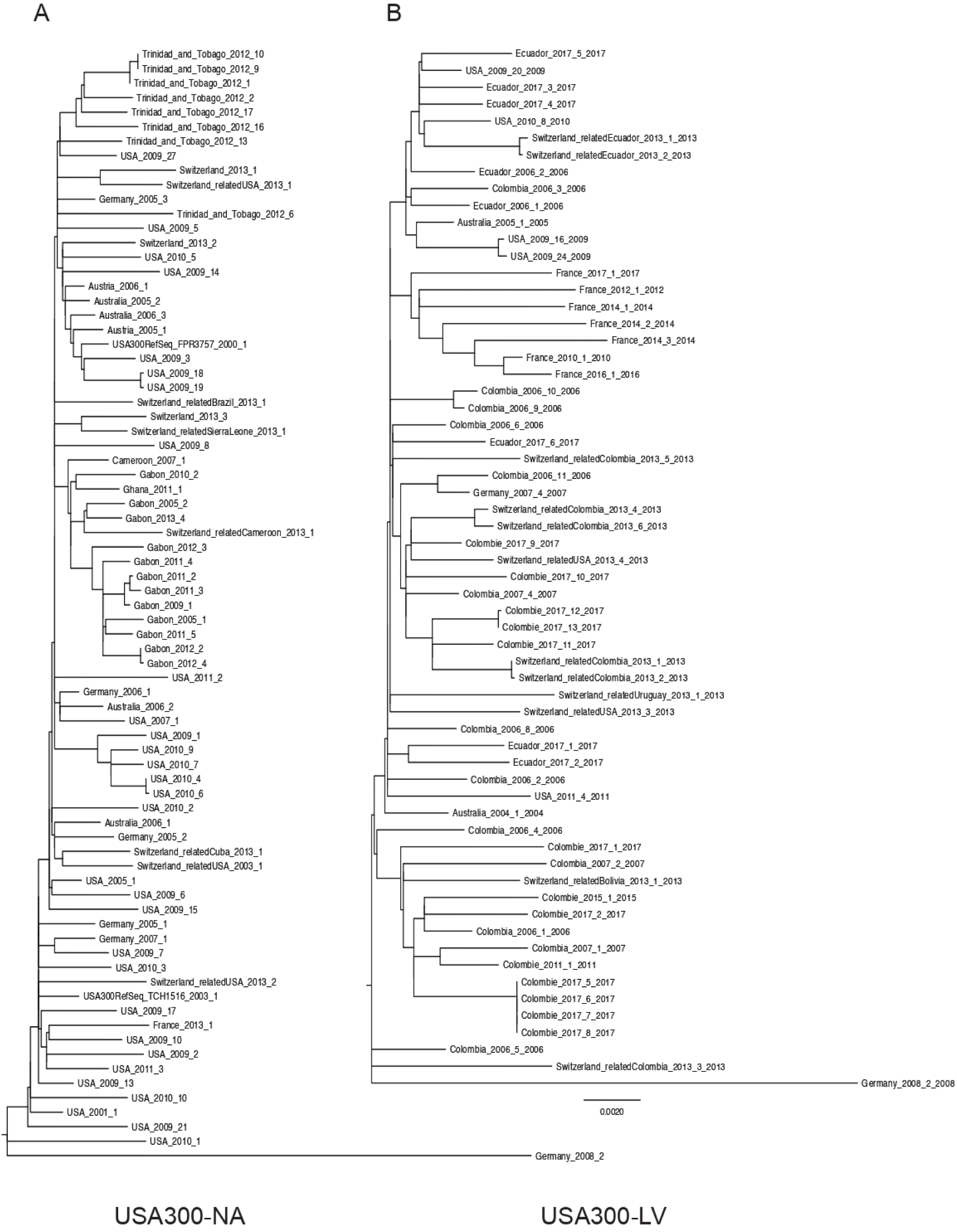
Evolutionary relationships of the selected strains belonging to the two USA300 sublineages. **(A)**, ML tree of the USA300-NA clade. **(B)**, ML tree of the USA300-LV clade. Note that both clades and rooted with the Germany_2008_2 strain. We implemented the general time reversible (GTR) + Γ model of DNA sequence evolution for both phylogenetic reconstructions.

**Figure S3.**
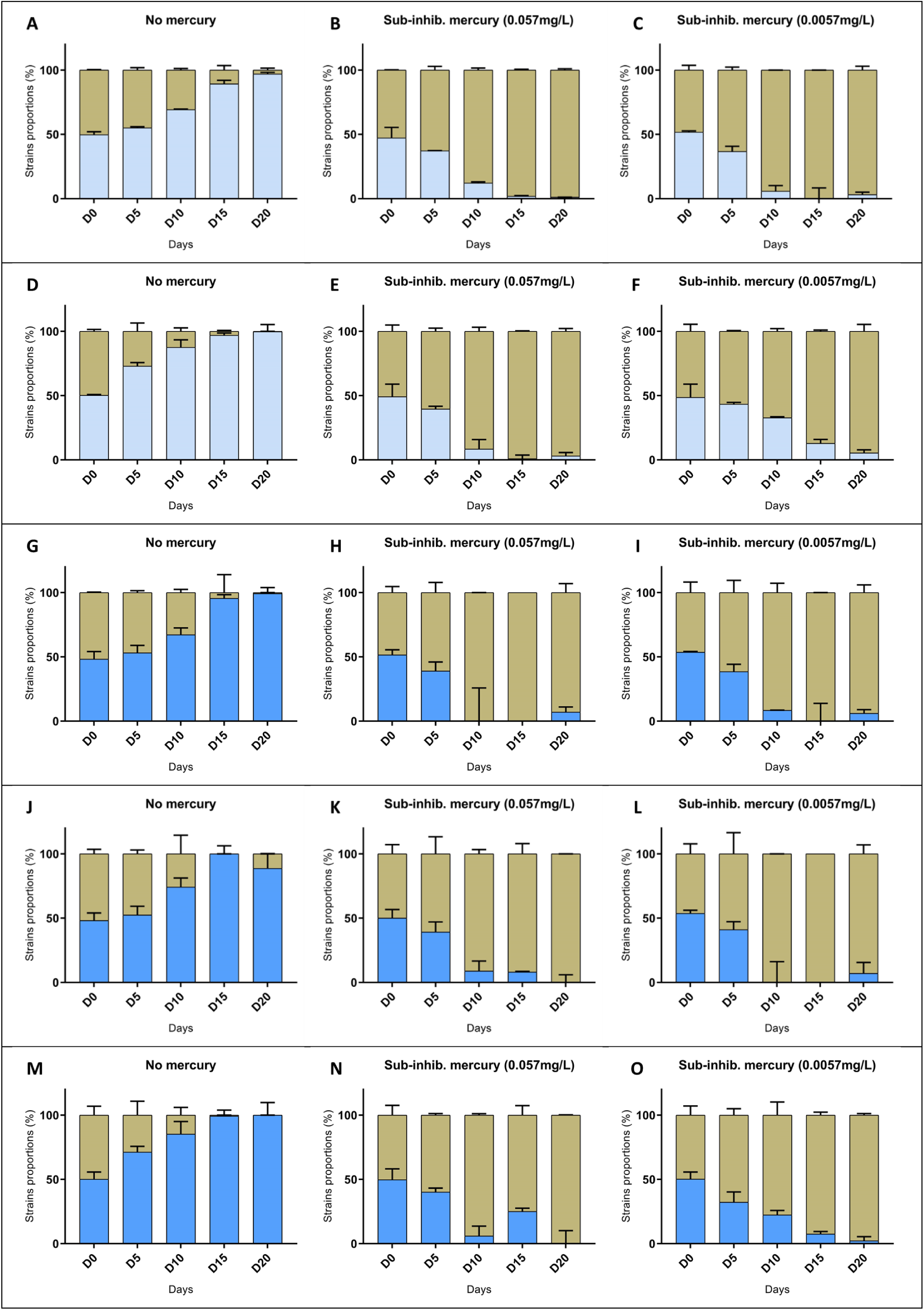
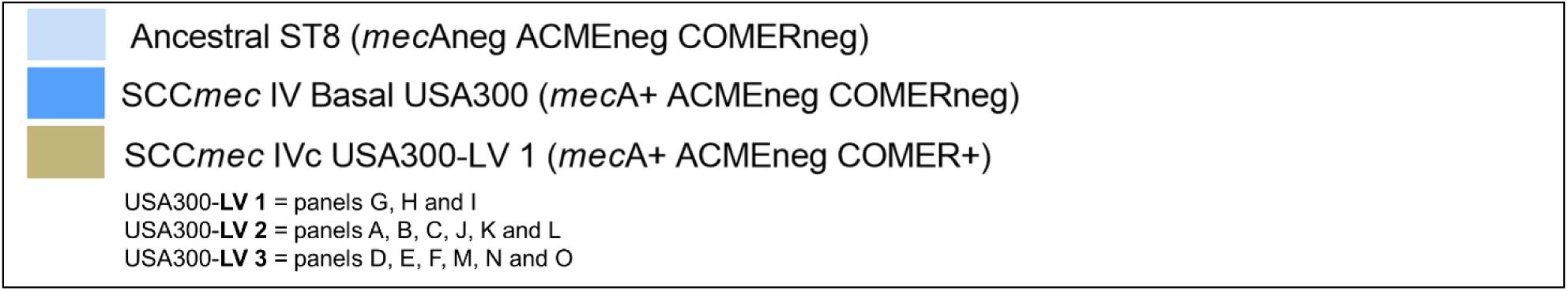
Impact of COMER and mercury resistance on competitive fitness of USA300 ancestral strains and LV variants. **(A) & (D)** USA300-LV (COMER-positive) strains and ancestral ST8 MSSA were co-cultivated for 20 days in BHI without antibiotics or **(B) & (E)** containing HgCl_2_ at ^1^/_10_ of HgCl_2_ MIC of the susceptible strain (0.057 mg/L), or **(C) & (F)** ^1^*/*_100_ MIC (0.0057 mg/L) with daily subculture in fresh medium. **(G), (J) & (M)** USA300-LV (COMER-positive) strains and Basal USA300 MRSA were co-cultivated for 20 days in BHI without antibiotics or **(H), (K) & (N)** containing HgCl_2_ at ^1^/_10_ of HgCl_2_ MIC of the susceptible strain (0.057 mg/L), or **(I), (L) & (O)** ^1^*/*_100_ MIC (0.0057 mg/L) with daily subculture in fresh medium. The proportion of each strain was monitored at day 0, 6, 10, 15 and 20 with differential colony count based on selective agar inoculated with a calibrated amount of competitive mix. Competitive cultures were performed on three independent series (biological replicates), and each colony count was repeated three times (technical replicates).

**Figure S4.**
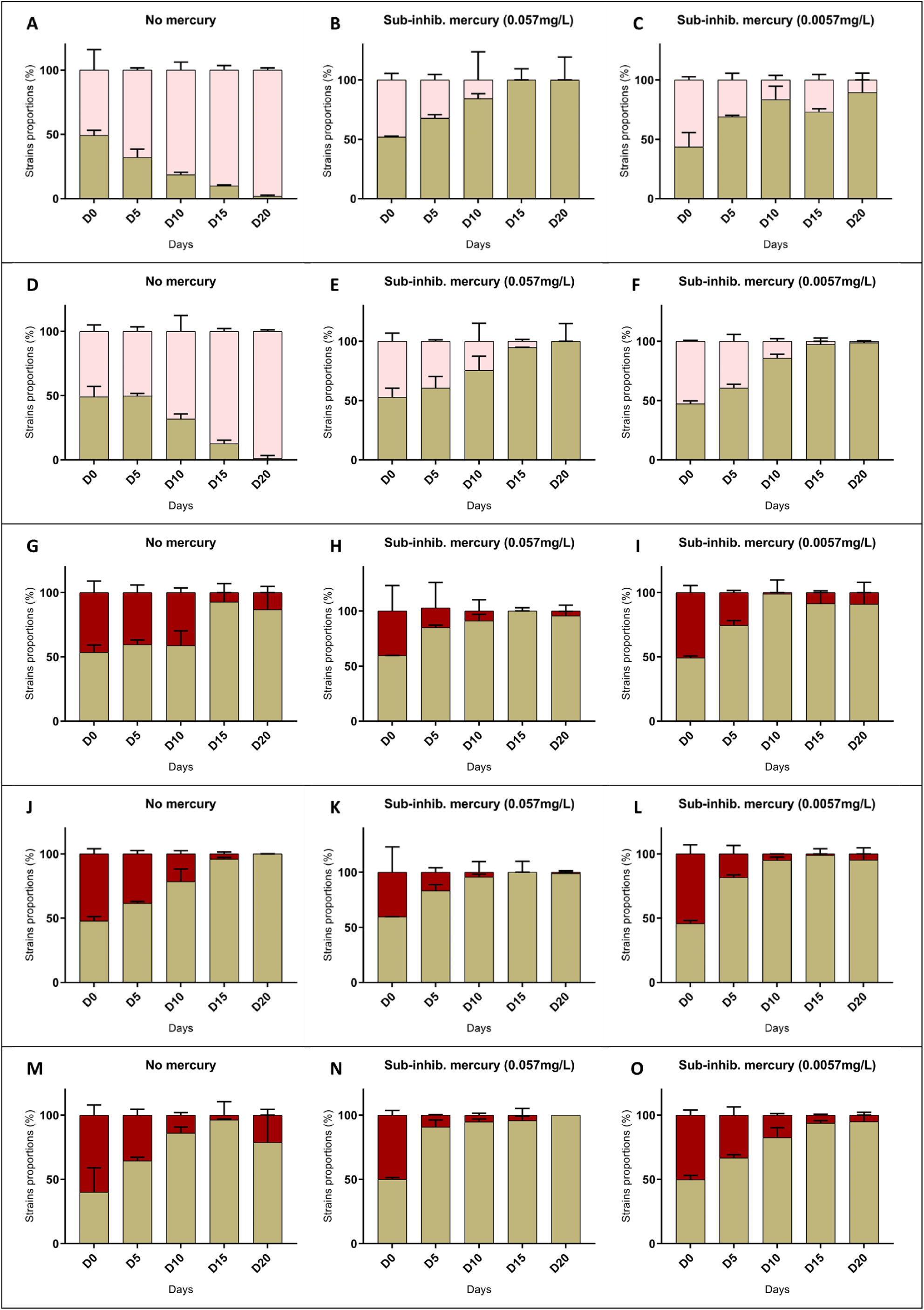
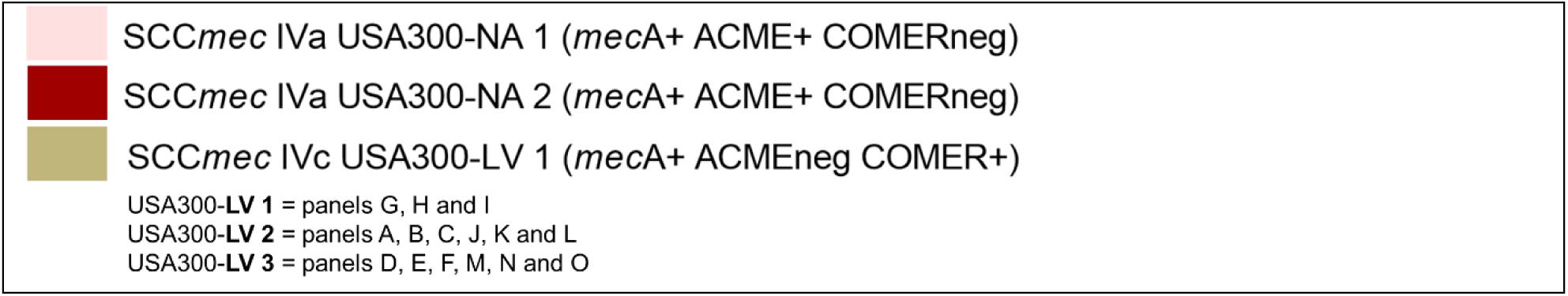
Impact of ACME, COMER and mercury resistance on competitive fitness of USA300 NA strains and LV variants. **(A) & (D)** USA300-LV (COMER-positive) strains and early derived USA300-NA (ACME-positive) were co-cultivated for 20 days in BHI without antibiotics or **(B) & (E)** containing HgCl_2_ at ^1^/_10_ of HgCl_2_ MIC of the susceptible strain (0.057 mg/L), or **(C) & (F)** ^1^*/*_100_ MIC (0.0057 mg/L) with daily subculture in fresh medium. **(G), (J) & (M)** USA300-LV (COMER-positive) strains and late derived USA300-NA (ACME-positive) were co-cultivated for 20 days in BHI without antibiotics or **(H), (K) & (N)** containing HgCl_2_ at ^1^/_10_ of HgCl_2_ MIC of the susceptible strain (0.057 mg/L), or **(I), (L) & (O)** ^1^*/*_100_ MIC (0.0057 mg/L) with daily subculture in fresh medium. The proportion of each strain was monitored at day 0, 6, 10, 15 and 20 with differential colony count based on selective agar inoculated with a calibrated amount of competitive mix. Competitive cultures were performed on three independent series (biological replicates), and each colony count was repeated three times (technical replicates).

